# Positive control of the integrative and conjugative element Tn*916* and its limited gene expression in transconjugants

**DOI:** 10.64898/2026.10.01.756105

**Authors:** Alexa F. S. Gomberg, Alan D. Grossman

## Abstract

Integrative and conjugative elements (ICEs) are widespread in bacteria and typically confer important phenotypes to their hosts. Following excision from the host chromosome, an ICE can transfer to a new cell through the element-encoded conjugation machinery (a type IV secretion system) to create a transconjugant. Tn*916*, a well-studied ICE that confers tetracycline resistance, is found in several Gram-positive pathogens and has been widely studied in *Bacillus subtilis*. Unlike most ICEs, Tn*916* causes host-cell death following its activation (excision and gene expression). We found that *orf7* and *orf8* (renamed *sig7* and *sig8*), two regulatory genes in Tn*916*, encode a two-subunit (split) sigma factor of RNA polymerase, Sig7-8, that activates transcription from P*orf7* (now P*sig7*) and is required for excision of the element. Importantly, production of Sig7-8, either from P*sig7* or another strong promoter, caused cell death. Using a Cre recombinase reporter to detect gene expression, we detected little or no transcription from P*sig7* in transconjugants. This lack of transcription from P*sig7* in transconjugants was essential for fitness of both the element and its host: forcing transcription from P*sig7* in transconjugants caused either loss of Tn*916* or death of nascent transconjugants. Our findings indicate that, unlike other well-studied ICEs (e.g., ICE*Bs1*) that use repressors to control gene expression and are expressed in nascent transconjugants, most Tn*916* genes remain off following transfer to a new host. This regulation prevents Tn*916* from killing transconjugants and allows successful horizontal transfer and maintains the potential for future activation in the new host.

## Introduction

Bacterial genomes can be modified by horizontal gene transfer (HGT). Conjugation, a type of HGT, requires cell-to-cell contact for conjugative plasmids and integrative and conjugative elements (ICEs) to transfer from a donor to a recipient. ICEs, a prevalent type of conjugative element (Guglielmini et al., 2011), are typically found integrated in the chromosome of a bacterial host and often confer beneficial traits to their host. In a subpopulation of cells, ICEs can excise from the chromosome and transfer to a neighboring cell through the element-encoded type IV secretion system (T4SS) (Johnson and Grossman, 2015). Genes required for element replication and transfer are often contiguous and regulated by a single promoter, and transcription from this promoter is typically inactive while the element is integrated. Inactivity is established either due to an element-encoded repressor or the absence of a functioning transcriptional activator (Auchtung et al., 2005; Beaber and Waldor, 2004).

Tn*916* is an ICE that confers tetracycline resistance to host cells (Roberts and Mullany, 2009). When integrated, the DNA processing and conjugation genes are not transcribed because they are separated from the main element promoter P*orf7* (Fig. 1A). Following excision and circularization, transcription from P*orf7* drives expression of the DNA processing and T4SS (conjugation) genes (Celli and Trieu-Cuot, 1998). P*orf7* is activated by the products of *orf7* and *orf8* and it has been proposed that it is repressed by the product of *orf9* (Celli and Trieu-Cuot, 1998; Roberts and Mullany, 2009). Current models propose that Tn*916* genes at the right end of the element (Fig. 1A) are initially expressed at low levels via readthrough transcription from P*tet*. In a subpopulation of cells, the amounts of Orf7 and Orf8 reach a level sufficient to activate transcription from P*orf7*, thereby driving robust expression of *xis* and *int*, causing excision and robust expression of all Tn*916* genes (Celli and Trieu-Cuot, 1998).

**Figure 1.**
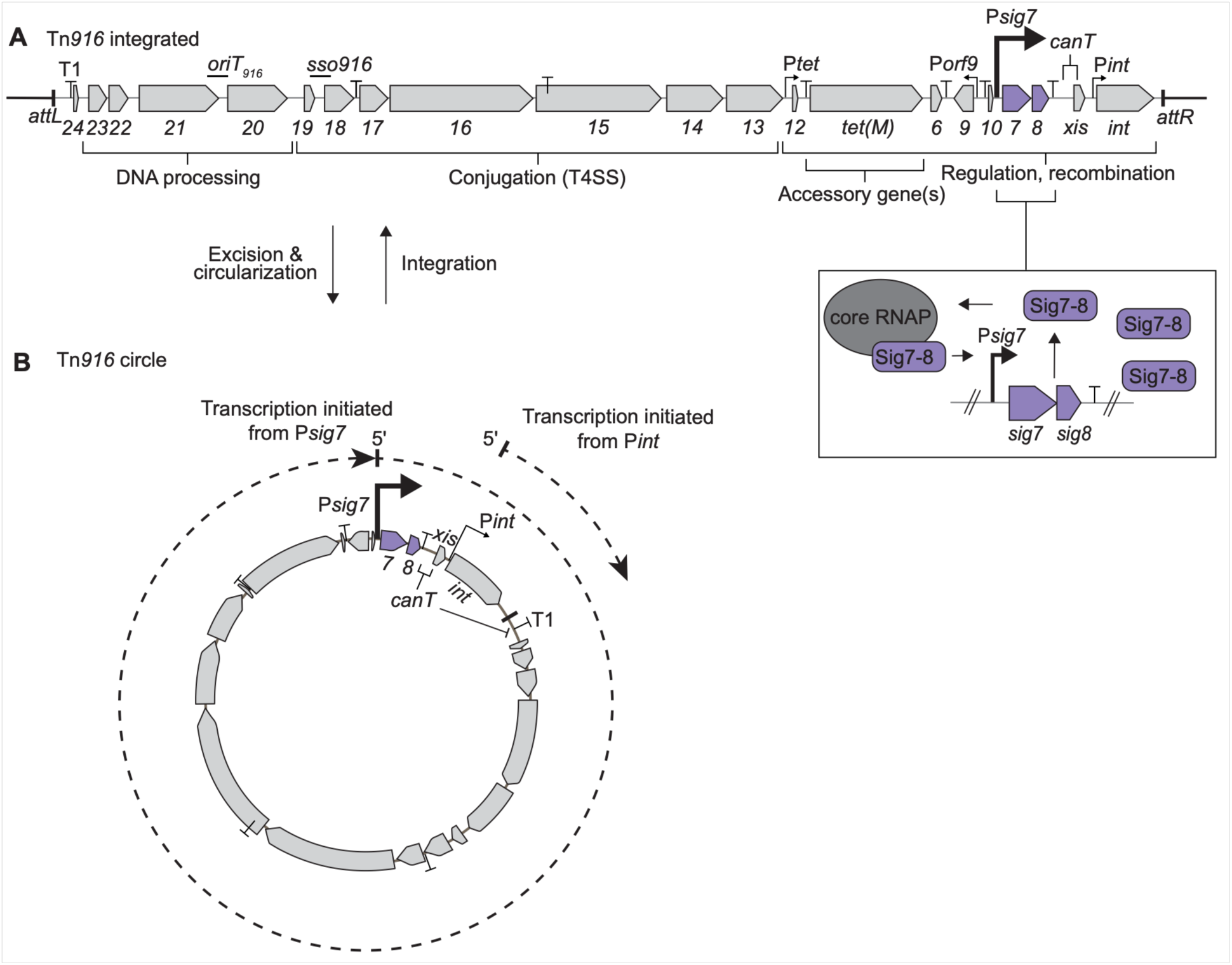
Maps of Tn*916*. Gene names (numbers) are indicated below the corresponding open reading frame. Promoters are indicated by bent arrows and transcription terminators are indicated by “T”s. Maps are adapted and modified from (Wirachman and Grossman, 2024). **A) Map of the integrated, linear Tn*916*.** The left and right ends of the integrated Tn*916* are indicated by vertical lines and *attL* and *attR*. The terminator T1 at the left end of the integrated element (Wirachman and Grossman, 2024), the origin of transfer (*oriT_916_*) (Jaworski and Clewell, 1995), the single strand origin of replication (*sso916*) (Wright and Grossman, 2016), and antiterminator (*canT*) (Wirachman and Grossman, 2024) are indicated above the map. Functional modules of genes are indicated by brackets below the map. The box on the right below the regulation module shows an expanded view highlighting the positive feedback loop caused by auto-activation of P*sig7* by Sig7-8. Sig7-8 (purple rectangles) associate with core RNAP (gray oval) to initiate transcription from P*sig7*. **B) The excised circular form of Tn*916*.** Following excision, P*sig7*, the main Tn*916* promoter, drives expression of the element genes (Celli and Trieu-Cuot, 1998). Transcripts from P*sig7* are not terminated at the T1 terminator due to the antiterminator *canT* (Wirachman and Grossman, 2024). Transcripts produced from P*int* do not include *canT* and are terminated at T1. *xis* is indicated above the map. *sig7-8*, *canT*, and *int* are indicated below the map.

Cells expressing the Tn*916* DNA processing and conjugation genes stop growing and die (Bean et al., 2022). This occurs both when these genes are expressed from a heterologous regulatory system in *Bacillus subtilis* and in cells with an activated Tn*916* (excised, circularized, and transcriptionally active) in *B. subtilis* and *Enterococcus faecalis* (Bean et al., 2022) and probably other host bacterial species. Multiple Tn*916* genes contribute to host cell death through at least three mechanisms (Bean et al., 2022) that differ substantially from those of two other ICEs, ICE*clc* (Reinhard et al., 2013; Takano et al., 2019) and Tn*Smu1* (McLellan et al., 2022) that also impair host cell growth and viability (Gomberg and Grossman, 2024). Despite the lethality caused by Tn*916* activation in donor cells, transconjugants survive the mating process. Little is known about how Tn*916* gene expression is regulated in nascent transconjugants (cells that have just received the element from a donor).

In this work, we investigated how Tn*916* is regulated, with emphasis on regulation in transconjugants. We found that *orf7* and *orf8* (now *sig7* and *sig8)* were essential for excision and conjugation and encode the major Tn*916* regulators. They encode a two-subunit (split) sigma factor, designated Sig7-8, that associates with RNA polymerase. Sig7-8 directs transcription from P*orf7* (P*sig7* below), which extends into *sig7-8* and the DNA processing and conjugation genes in the circular form of Tn*916*. We found that expression of *sig7-8* alone, in the absence of other Tn*916* genes, was sufficient to cause cell death. We also found that there was little or no expression from P*sig7* in transconjugants. Hence, *sig7-8* and the conjugation genes that cause cell death are not appreciably expressed in transconjugants, thereby enabling their survival. In contrast, and as expected, there was transcription from Tn*916* P*int* in transconjugants. P*int* drives expression of *int* which encodes the recombinase needed for element integration and excision. Together, our results provide an integrated view of how Tn*916* causes cell death and how cells that acquire Tn*916* via conjugation survive while the donor cells do not.

## Results

### Rationale

Following excision and circularization of Tn*916* (Fig. 1B), P*orf7* is the main promoter that drives expression of the Tn*916* genes needed for conjugation (Celli and Trieu-Cuot, 1998). Analyses of promoters cloned from Tn*916* and fused to *lacZ* demonstrated that both *orf7* and *orf8* are needed for transcription from P*orf7* (Celli and Trieu-Cuot, 1998). It is not known how *orfs7-8* function in activating transcription from P*orf7*, nor what effects their loss has on Tn*916* excision and conjugation in an otherwise intact element.

As previously recognized (Su and Clewell, 1993), Orf7 is similar to the C-terminal region (region 4) of sigma factors of RNA polymerase. Sigma factors associate with core RNA polymerase to form a holoenzyme that recognizes promoters and initiates transcription (Feklístov et al., 2014; Helmann, 2019; Helmann and Moran Jr., 2001). Canonical sigma-70-like factors have four domains (regions 1-4) in a single protein. In contrast, some alternative sigma factors (group 4) contain only regions 2 and 4 (Helmann, 2002; Lonetto et al., 1992), typically in a single protein. Region 4 of sigma factors interacts with the -35 region of promoters (Lonetto et al., 1992). Orf8 was not predicted to be a sigma factor. SeqHub, a platform that uses a protein language model system based on Gaia (Jha et al., 2025), indicated that Sig8 is similar to RsoA, a component of a two-subunit split sigma factor. Together, the proteins in split sigma factors contain sigma factor regions 2 and 4 and the two proteins associate with RNA polymerase and recognize the -10 and -35 regions of promoters (Helmann, 2002; Lonetto et al., 1992; MacLellan et al., 2008). Based on these similarities, we investigated the function of Orf7 and Orf8 and experiments presented below indicate that Orf7 and Orf8 (now Sig7 and Sig8) associate with RNA polymerase and are two subunits of a split sigma factor.

### Sig7 and Sig8 associate with RNA polymerase

Sigma factors associate with and typically can be isolated with RNA polymerase. We used a C-terminal histidine-tagged β’ subunit of RNA polymerase (β’-His) (Qi and Hulett, 1998) to enable simple isolation of the enzyme and associated factors. To monitor association of Sig7 and Sig8 with RNA polymerase, we tagged the C-terminus of each with either a FLAG (Sig7-FLAG) or a Myc epitope (Sig8-Myc) (Methods). Both were expressed from the strong inducible promoter P*xis* from ICE*Bs1* (Fig. 2A; Methods).

**Figure 2.**
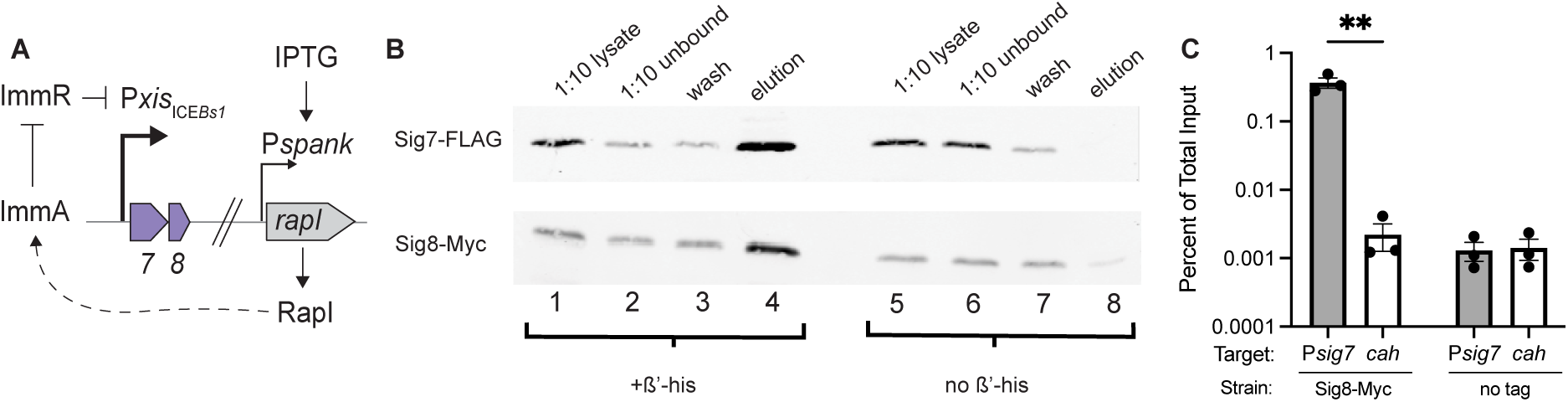
Sig7 and Sig8 are a two-subunit (split) sigma factor. **A)** Schematic of regulation of P*xis*, the promoter from ICE*Bs1* that was used for ectopic expression of *sig7-8*. The controlled de-repression of P*xis* has been described (Bose et al., 2008). Briefly, P*xis* is repressed by ImmR, ImmR is cleaved by the protease ImmA, and the reaction is stimulated by RapI. In this ectopic system, *rapI* is expressed from the LacI-repressible IPTG-inducible promoter Pspank integrated in the chromosome at *lacA* and *immR* and *immA* are expressed from their own promoter (P*immR*) and integrated in the chromosome at *amyE* with P*xis* and *sig7-8* (Table 1). Addition of IPTG causes de-repression of P*xis* and production of Sig7-8. **B)** Western blot probed with anti-FLAG (top) or anti-Myc (bottom) antibodies to detect Sig7-FLAG and Sig8-Myc, respectively. Lanes 1-4: samples collected from cells expressing *rpoC-his10* [+ß’-His; strain AFG723]. Lanes 5-8: samples collected from cells without the *his-*tag [No ß’-His; strain AFG728]. Lanes 1, 5: 1:10 dilution of crude cell lysate before affinity purification (on cobalt resin) of RNA polymerase and associated proteins. Lanes 2, 6: 1:10 dilution of the unbound fraction (that did not associate with cobalt resin). Lanes 3, 7: undiluted first wash of the cobalt resin. Lanes 4, 8: undiluted elution from the cobalt resin. Data are from one representative experiment. Similar results were obtained in three independent experiments. **C)** Association of Sig8-Myc with P*sig7*. *sig8-myc* was expressed from P*xis* (as above). P*sig7*-*mapple2* was integrated in the chromosome at *ycgO*. Left two bars: Sig8-Myc [strain AFG720]; Right two bars: a specificity control that lacked the Myc tag [no tag; strain AFG546]. Cells were grown, treated with formaldehyde to crosslink, and samples were immunoprecipitated with anti-Myc antibodies (Methods). DNA in the immunoprecipitates was detected by qPCR with primers specific to P*sig7* (filled bars) and the nearby chromosomal locus *cah* (∼5 kb from P*sig7*) (open bars). Percentage of total input (y-axis) was calculated as the ratio of the amount of DNA from the indicated region (as determined by qPCR) following immunoprecipitation compared with that of the indicated region prior to immunoprecipitation, times 100%. Data are presented from three independent experiments with error bars depicting standard error of the mean (mean ± SEM). P-values were calculated by a multiple t-test comparison followed by Bonferroni multiple test correction (**P < 0.01) using the GraphPad Prism version 11. The two samples in the specificity control with no tag were indistinguishable. Attempts to detect P*sig7* in the same crosslinking reactions by immunoprecipitating Sig7-FLAG were unsuccessful. We do not know if this is due to lack of crosslinks between Sig7 and the DNA or inefficient precipitation of the FLAG epitope.

**Table 1.**
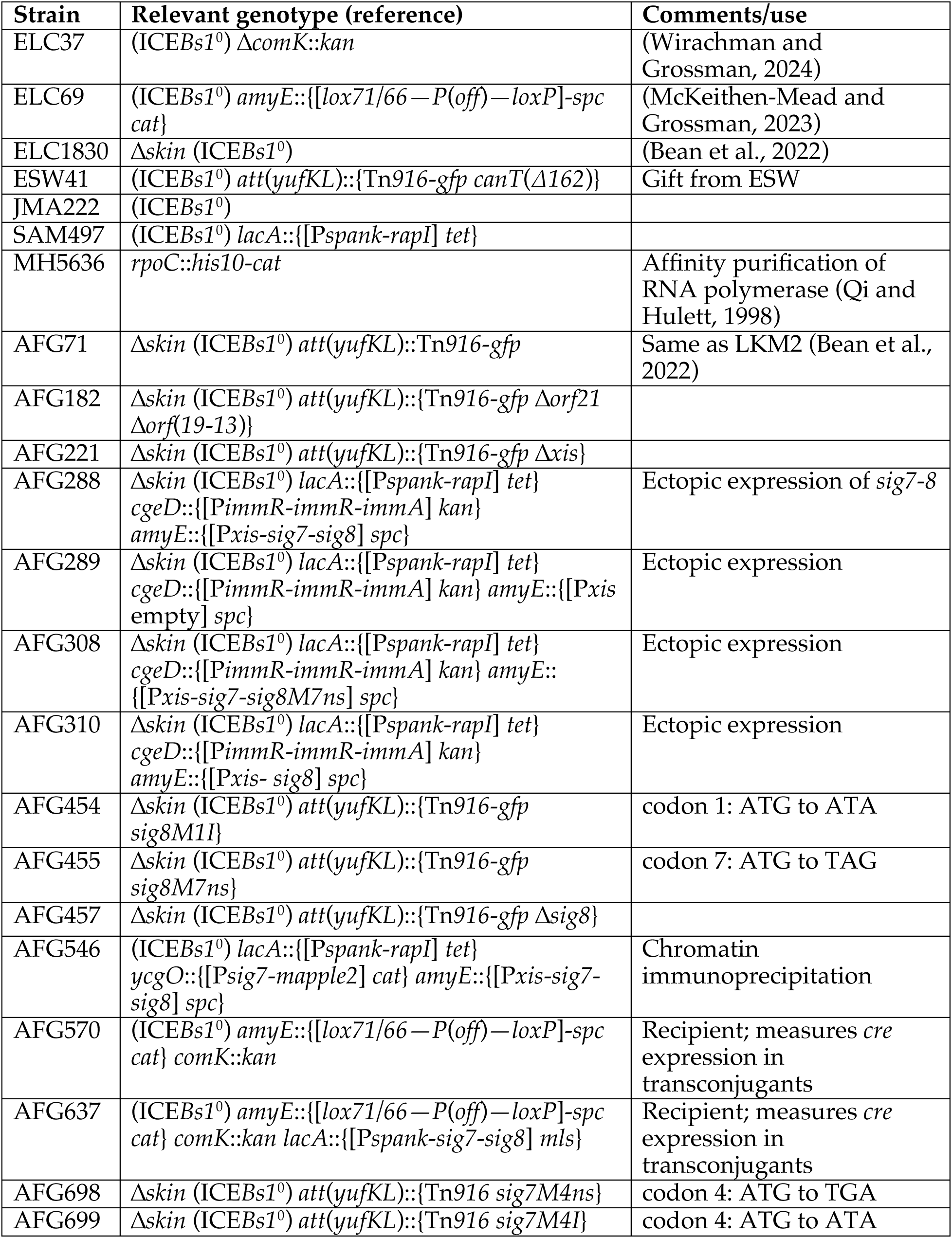
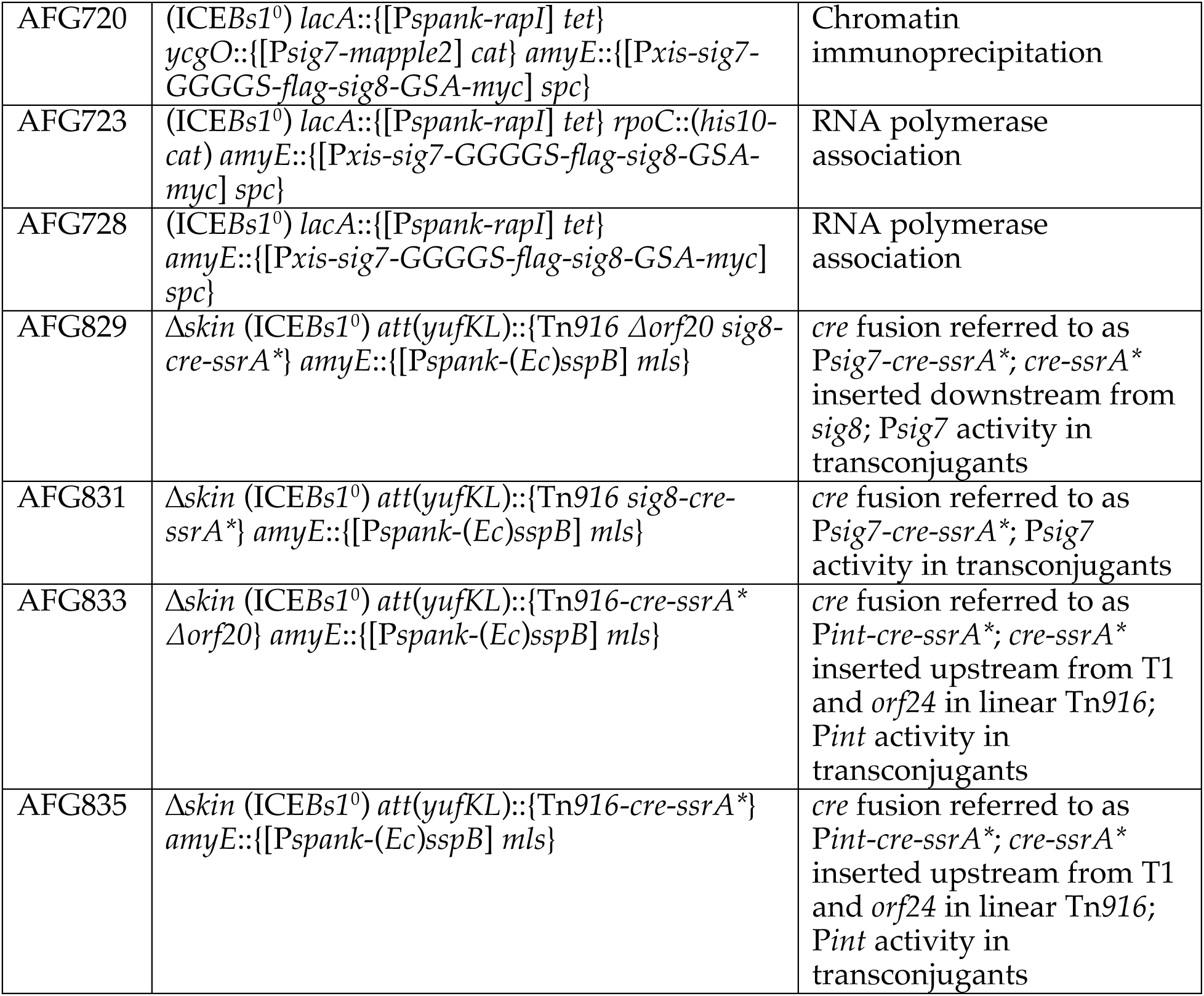
*B. subtilis* strains used.

We found that Sig7 and Sig8 associated with *B. subtilis* RNA polymerase (Fig. 2B). Cells were grown in defined minimal medium and expression of *sig7-flag* and *sig8-myc* was induced for one hour with IPTG (Methods). We isolated the His-tagged RNA polymerase on cobalt resin and detected Sig7-FLAG and Sig8-Myc after elution of the RNA polymerase (Methods). Using Western blotting and antibodies to each epitope, we detected both Sig7-FLAG (Fig. 2B; top panels) and Sig8-Myc (Fig. 2B; bottom panels) in the crude lysate before purification (Fig. 2B; lanes 1, 5), fractions that did not bind to the resin (lanes 2, 6), and washes (lanes 3, 7). Importantly, both Sig7-FLAG and Sig8-Myc were associated with RNA polymerase and eluted from the resin (Fig. 2B; lane 4). This association required the presence of the His-tagged RNA polymerase. In the absence of β’-His (i.e., with untagged RNA polymerase), there was little or no detectable association of Sig7-FLAG or Sig8-Myc with the cobalt resin (Fig. 2B; lane 8).

Based on the association of both Sig7 and Sig8 with RNA polymerase and their sequence similarity to sigma factors, we conclude that Sig7 and Sig8 from Tn*916* most likely comprise a two-subunit (split) sigma factor. Because they are both needed for transcription from P*sig7* (Celli and Trieu-Cuot, 1998), they are most likely directing RNA polymerase to P*sig7* of Tn*916*.

#### Association with P*sig7*

Sigma factors associate with RNA polymerase to recognize and bind cognate promoters. We tested directly for association of Sig8-Myc with P*sig7* using chromatin immunoprecipitation. We crosslinked protein and DNA with formaldehyde (Methods) in cells expressing *sig8-myc* and *sig7-flag*, immunoprecipitated Sig8-Myc, and tested for co-precipitation of DNA from P*sig7*. There was >100-fold enrichment of P*sig7* DNA in the immunoprecipitation compared to that of a nearby locus, *cah* (∼5 kb away) (Fig. 2C). Based on these results, together with data in the literature on gene expression, we conclude that transcription from P*sig7* is driven by RNA polymerase containing the two-subunit sigma factor Sig7-8 and that this transcription drives expression of Tn*916* genes following excision and circularization of the element. In addition, transcription from P*sig7* creates a positive-feedback loop further increasing production of Sig7 and Sig8 (Fig. 1A).

### *sig7* and *sig8* are needed for excision and conjugation of Tn*916*

Previous work demonstrated that *orf7* and *orf8* (now *sig7* and *sig8*) are required for transcription from P*orf7* (now P*sig7*) in a reporter construct (Celli and Trieu-Cuot, 1998). It was proposed that these are required for both excision and conjugation (Celli and Trieu-Cuot, 1998; Hinerfeld and Churchward, 2001; Marra et al., 1999; Roberts and Mullany, 2009), the implication being that the promoter upstream of *xis* was not sufficient for excision (Celli and Trieu-Cuot, 1998; Marra et al., 1999). To investigate the functional requirement for *sig7* and *sig8* in the context of an intact Tn*916*, we generated nonsense mutations in both *sig7* and *sig8*. However, mutagenesis of *sig7* first revealed that the annotated start codon is likely not the true translation initiation site.

Two possible products for *sig7* (then *orf7*) were initially proposed (Su and Clewell, 1993). The predicted open reading frames were referred to as *orf7.1* and *orf7.2* (S1A Fig), with predicted start codons ATA and ATG, respectively (Su and Clewell, 1993). However, the predicted initiation codon for *orf7.1* (ATA) is rarely used as an initiation codon (Hecht et al., 2017; Kohl et al., 2026; Peijnenburg et al., 1990; Romero and García, 1991). Preliminary mutational analyses indicated that the GTG that is three codons upstream from the ATG of the predicted *orf7.2* is likely the start codon (S1 Fig). For analyses below, we designate the GTG to be the start, and the ATG to be the fourth codon (S1A Fig). In this case, Sig7 would be 143 amino acids.

We made nonsense mutations in *sig7* and *sig8*. We changed the ATG (labeled codon 4) to the nonsense codon TGA (*sig7M4ns*) (S1A Fig) and changed codon 7 (ATG) of *sig8* to the nonsense codon TAG (*sig8M7ns*). Each mutation was introduced separately into Tn*916* and their effects on excision and conjugation frequencies were determined. Excision was measured using qPCR to detect the vacant chromosomal site created following Tn*916* excision (Methods). Wild-type Tn*916* excised at a frequency of ∼10^−2^ per cell (Fig. 3A). In contrast, the *sig7* and *sig8* mutants had little or no detectable excision, with the qPCR signal similar to that of an *xis* null mutant that is incapable of excision (Fig. 3A).

**Figure 3.**
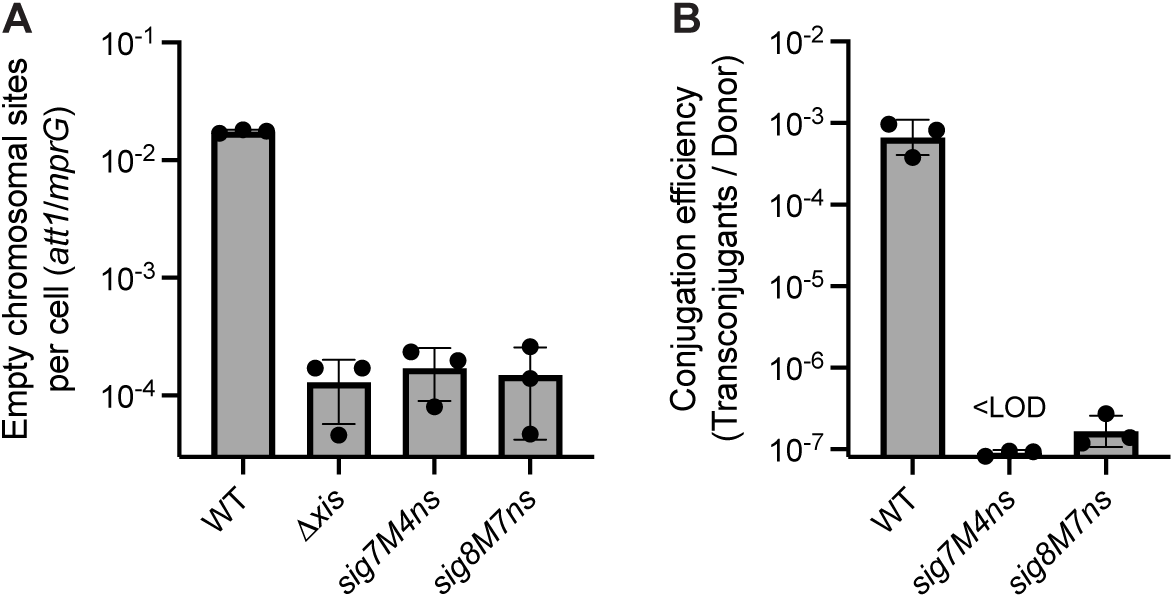
*sig7-8* are needed for mating and excision. Excision (A) and mating efficiencies (B) of wild-type Tn*916* [WT, AFG71], Tn*916* with a nonsense mutation in *sig7* [*sig7*(*M4ns*); strain AFG698], and Tn*916* with a nonsense mutation in *sig8* [*sig8*(*M17ns*); strain AFG455]. **A)** Excision frequency was determined by measuring the amount of DNA from an ’empty’ integration site (*yufKL*), indicative of Tn*916* excision compared to the amount of DNA from a nearby site (*mrpG*). Tn*916 Δxis* [strain AFG221] is incapable of excision and was used as a control to determine the limit of detection in the qPCR assay, and all measurements of excision of the *sig7*(*M4ns*) and *sig8*(*M7ns*) mutants were indistinguishable from that of the *Δxis* mutant. **B)** Conjugation efficiencies were calculated from mating the indicated donor (above and indicated below each bar) with a kanamycin-resistant recipient [strain ELC37], and measuring the number of transconjugants (resistant to tetracycline and kanamycin) per initial donor. The mating efficiency of the *sig7* mutant was at or below the limit of detection. In one mating assay of the *sig8* mutant, there were apparently three transconjugants (just above the limit of detection). There were no detectable transconjugants in other independent experiments with the *sig8* mutant.

As expected from the lack of excision, the mutants were also defective in conjugation. Wild-type Tn*916* had a conjugation efficiency of ∼10^−3^ transconjugants per donor (Fig. 3B). In contrast, conjugation of the *sig7* and *sig8* mutants was at or below the limit of detection in these experiments (< 2 × 10^−7^ transconjugants per donor) (Fig. 3B). These results demonstrate that, as previously proposed, *sig7* and *sig8* are required for detectable excision and conjugation of Tn*916* under the conditions tested.

### Analysis of Tn*916* mutants for effects on cell growth

Most cells with an activated Tn*916* (excised circular form with genes expressed) stop growing and die (Bean et al., 2022). There are two previously identified mechanisms contributing to growth arrest and cell death: one mechanism is mediated by *yqaR* (found in the defective phage *skin*) and *orfs17-16*; a second mechanism is mediated by *orfs 21, 19,* and *15* and is independent of *yqaR* (Bean et al., 2022). We wished to determine if Sig7-8 contributed to the effect on cell growth, separate from its role in transcription of Tn*916* conjugation genes. Experiments described below demonstrate that high levels of Sig7-8 also cause growth arrest and cell death, both in the context of Tn*916* and when they are expressed ectopically, without any other Tn*916* genes.

We used a Tn*916*-*gfp* to track cells in which the element had excised and circularized, essentially as described previously (Bean et al., 2022). Briefly, *gfp* is very near the left end of Tn*916*, between the integration site and the transcription terminator T1 (Fig. 4A). There is no promoter driving expression of *gfp* until the element excises. After excision (activation), *gfp* is downstream of and transcribed from P*sig7* and activated cells can be identified by green fluorescence. Expression of genes in the circular form of Tn*916* is dependent on the transcription antiterminator *canT*, although expression of *gfp* is not because it is encoded upstream of the terminator T1 (Wirachman and Grossman, 2024).

**Figure 4.**
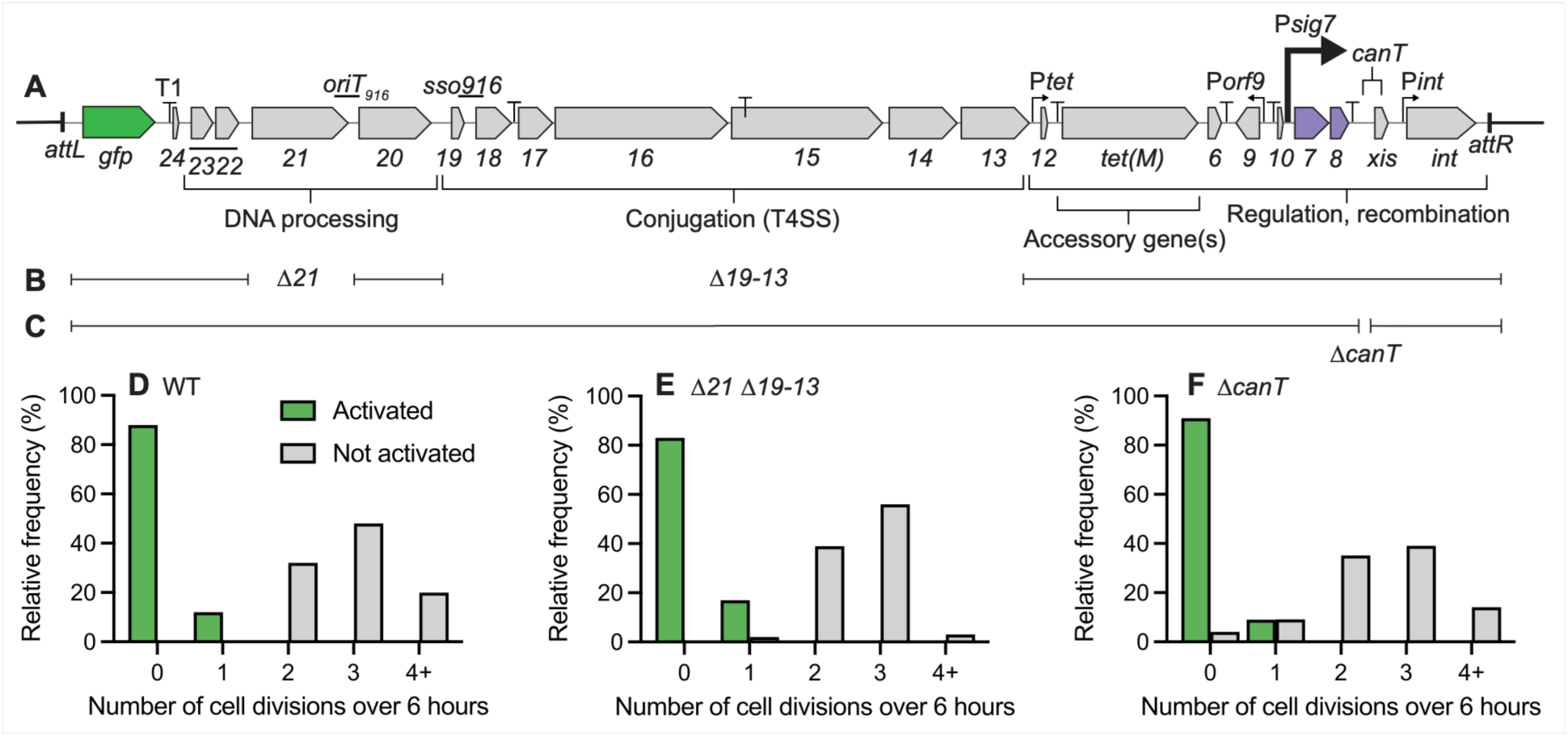
Effects of Tn*916* mutations on host cell growth. **A-C.** Schematic of Tn*916*-*gfp* and the mutants analyzed. **A.** *gfp* is inserted at the left end of the integrated Tn*916*, upstream from the terminator T1. There is no promoter driving transcription of *gfp* until Tn*916* excises as a circle (Bean et al., 2022). **B.** Deletions *Δ21* and Δ(*19-13*) remove the gene encoding the coupling protein (*orf21*) and genes encoding the conjugation proteins *orfs*(*19-13*), and together these remove the genes previously found to contribute to arrest of host growth and cell death (Bean et al., 2022). **C.** *ΔcanT* removes the anti-terminator that is essential for transcription through the terminator T1 and expression of all genes downstream from T1 (Wirachman and Grossman, 2024). *ΔcanT* leaves the flanking genes *sig8* and *xis* intact. **D-F.** Growth arrest of cells with an activated Tn*916*. The indicated strains were grown and individual cells visualized by fluorescence microscopy. Cells with an activated Tn*916* were green and indicated by green bars. Cells without an activated Tn*916* did not fluoresce green and are indicated in gray bars. Cell growth and division was tracked for six hours and the percent of cells (y-axis) that underwent the indicated number of cell divisions (x-axis) after six hours is shown. **D.** WT; Wild type Tn*916-gfp* [strain AFG71]; 42 cells with an activated Tn*916* and 50 cells in which Tn*916* had not activated were tracked. **E.** *Δ21 Δ*(*19-13*); Tn*916-gfp* Δ*orf21* Δ*orf*(*19-13*) [strain AFG182]; 41 cells with an activated Tn*916* and 62 cells in which Tn*916* had not activated were tracked. **F.** *ΔcanT*; Tn*916-gfp canT*(*Δ162*) [strain ESW41]; 43 cells with an activated Tn*916* and 57 cells in which Tn*916* had not activated were tracked. Transcripts from Tn*916-gfp canT*(*Δ162*) are terminated at T1 and do not transcribe *orfs24-13*.

We deleted the five genes in Tn*916* that were previously identified that contributed to growth arrest and cell death (Bean et al., 2022). These genes included (gene products and Vir Agrobacterium homologs in parentheses) *orfs*: *21* (coupling protein; VirD4); *19* (membrane protein involved in conjugation with no homolog in Vir system); *17-16* (VirB3-like; VirB4); and *15* (VirB6-like). We constructed Tn*916-gfp* mutants that were missing these genes by deleting *orf21* and *orfs 19-13* (Fig. 4B). In addition, we used a mutant that is defective in the transcription antiterminator *canT* (Fig. 4C). This mutant has a normal excision frequency but has little to no expression of the conjugation genes and has no detectable conjugation (Wirachman and Grossman, 2024).

We found that all Tn*916* mutants tested still caused growth arrest in cells with an active element. Cells were grown in defined minimal medium and activation of Tn*916* was stimulated by addition of tetracycline. Fluorescence microscopy was used to track cells expressing *gfp*. In all the mutants tested, all tracked cells with an activated Tn*916-gfp* stopped growing while surrounding cells continued to grow and divide (Fig. 4D-F). This growth arrest was seen in the deletion mutants missing all known killing genes and in the *canT* mutant that should have little or no expression of genes downstream from the terminator T1.

All constructs tested expressed genes from *tetM* through the cloned *gfp* upstream of T1 (Fig. 4A). We therefore infer that there is at least one gene in this part of Tn*916* that causes growth arrest. Because *sig7-8* encode an alternate sigma factor that auto-activates, and some other alternate sigma factors can cause cell death when auto-activated or overexpressed (Burton et al., 2023; Schofield et al., 2021), we postulated that expression of *sig7-8* might cause growth arrest and cell death.

### Ectopic expression of *sig7* and *sig8*, in the absence of other Tn*916* genes, causes growth arrest and cell death

We found that ectopic expression of *sig7* and *sig8* from the strong ICE*Bs1* promoter P*xis* (P*xis*-*sig7-8*; integrated in single copy in the *B. subtilis* chromosome at *amyE*) caused host cell death. Cells were grown in defined minimal medium and IPTG was added to induce P*spank*-*rapI*. RapI causes cleavage of the repressor ImmR resulting in derepression of P*xis*-*sig7-8*. Cell growth slowed approximately two hours after expression of P*xis*-*sig7-8* (Fig. 5A). In contrast, cells with just the induction system and no *sig7-8* continued to grow exponentially after addition of IPTG (Fig. 5A).

**Figure 5.**
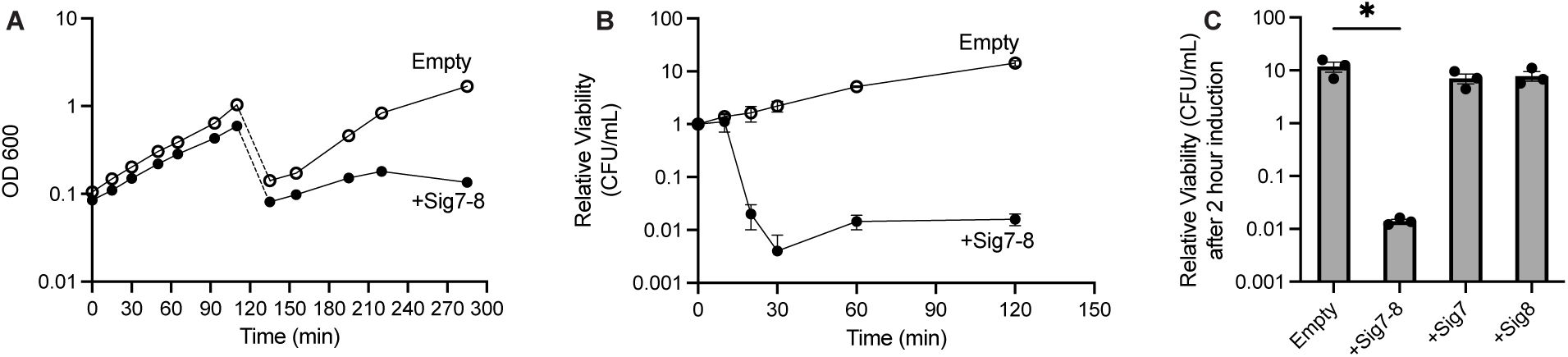
Ectopic expression of Sig7 and Sig8 causes host cell death. Strains with the inducible expression system P*xis* (Fig. 2A) driving transcription of *sig7-8* (+Sig7-8), *sig7* alone (+Sig7), *sig8* alone (+Sig8), or an empty vector (Empty), were grown to an OD600 of approximately 0.1 and expression from P*xis* was induced by addition of IPTG (1 mM). **A-B.** Empty (open circles) indicates cells with the P*xis* vector without an insert [strain AFG289]. +Sig7-8 (filled circles) indicates cells expressing P*xis*-*sig7-8* [strain AFG288]. **A.** Growth in cell culture was tracked by optical density (OD600) as a function of time. Dotted lines at about 120 minutes indicates dilution of cells 1:10 into fresh medium (with IPTG) to allow for more growth before entry into stationary phase. Data are presented from a representative of at least three independent experiments. **B.** Cell viability was measured as colony forming units (CFUs) at the indicated times after induction of P*xis* (addition of IPTG). CFUs at the indicated times were normalized to the CFUs of that culture at time 0 (before addition of IPTG). Data presented are averages from two independent experiments (including data from the experiment presented in panel A) with error bars depicting standard error of the mean (mean ± SEM). **C.** Sig7 and Sig8 are both required for loss of viability. CFUs were measured and compared to initial CFUs after 2 hours of induction with 1mM IPTG. Sig7-containing cells [+Sig7; strain AFG308] are isogenic to Sig7-8-containing cells but have a nonsense mutation in *sig8* (*sig8M7ns*). Sig8-containing cells [+Sig8; strain AFG310] contain only the coding sequence for *sig8*. Data presented are averages from three independent experiments with error bars depicting standard error of the mean (mean ± SEM). P-values were calculated by an ordinary one-way ANOVA comparison followed by Dunnett’s multiple test correction using the GraphPad Prism version 11. Significance comparisons were made against the empty vector control [Empty, AFG289]. * indicates P < 0.05. +Sig7 and +Sig8 were not significantly different from the empty vector control. Data from panels A and B are included here for the empty vector and cells expressing *sig7-8*.

We also found that cells expressing P*xis*-*sig7-8* lost viability, even before there was a notable effect on growth. We plated samples from liquid culture onto LB agar (without IPTG) at various times after expression of P*xis*-*sig7-8*. Cell viability dropped approximately 100-fold within 20 minutes after expression of P*xis*-*sig7-8* (Fig. 5B). Both *sig7* and *sig8* were needed for the drop in viability as expression of one or the other alone had little or no effect after two hours of induction (Fig. 5C). These results indicate that expression of the alternate sigma factor Sig7-8, in the absence of any other Tn*916* genes, is toxic to *B. subtilis*.

### Reporters to monitor transcription from Tn*916* promoters in nascent transconjugants

A majority of host cells with an active Tn*916* stop growing and die due to a combination of high levels of expression of *sig7-8* (above) and expression of some of the conjugation genes from P*sig7* (Bean et al., 2022). Yet transconjugants survive. We postulated that there might not be transcription from P*sig7* in nascent transconjugants, thereby limiting expression of *sig7-8* and the conjugation genes and allowing nascent transconjugants to remain viable. We also reasoned that because Tn*916* integrates into the chromosome of a new host, the recombinase gene *int* is most likely expressed from its promoter P*int* in the nascent transconjugant. We designed reporters to monitor expression from P*int* and P*sig7* in nascent transconjugants.

We inserted *cre* downstream of P*int* (Fig. 6A) or P*sig7* (Fig. 6B) in Tn*916* to measure transcription in nascent transconjugants. The reporter for the Cre recombinase was previously described and has an inverted promoter flanked by *lox* sites upstream from *spc* [Fig. 6C; (McKeithen-Mead and Grossman, 2023)]. Recipients with this reporter are sensitive to spectinomycin. Cre-mediated inversion between the *lox* sites results in an active promoter now driving transcription of *spc* and conferring spectinomycin resistance to viable cells (Fig. 6D). In addition, we can detect inversions by qPCR using primers that target the inverted region (Fig. 6D; primers a and b). Because transcription from P*sig7* is high in activated cells, we were concerned that abundant production of Cre could damage the host and/or transfer to recipients. To reduce accumulation of Cre, we made a *cre-ssrA\** fusion and expressed the adaptor protein SspB (from *E. coli*) in donors. The presence of SspB causes Clp-mediated degradation of SsrA*-tagged proteins (Griffith and Grossman, 2008). In this way, Cre-SsrA* would be selectively degraded in donors (in the presence of SspB) and more stable in recipients in the absence of SspB.

**Figure 6.**
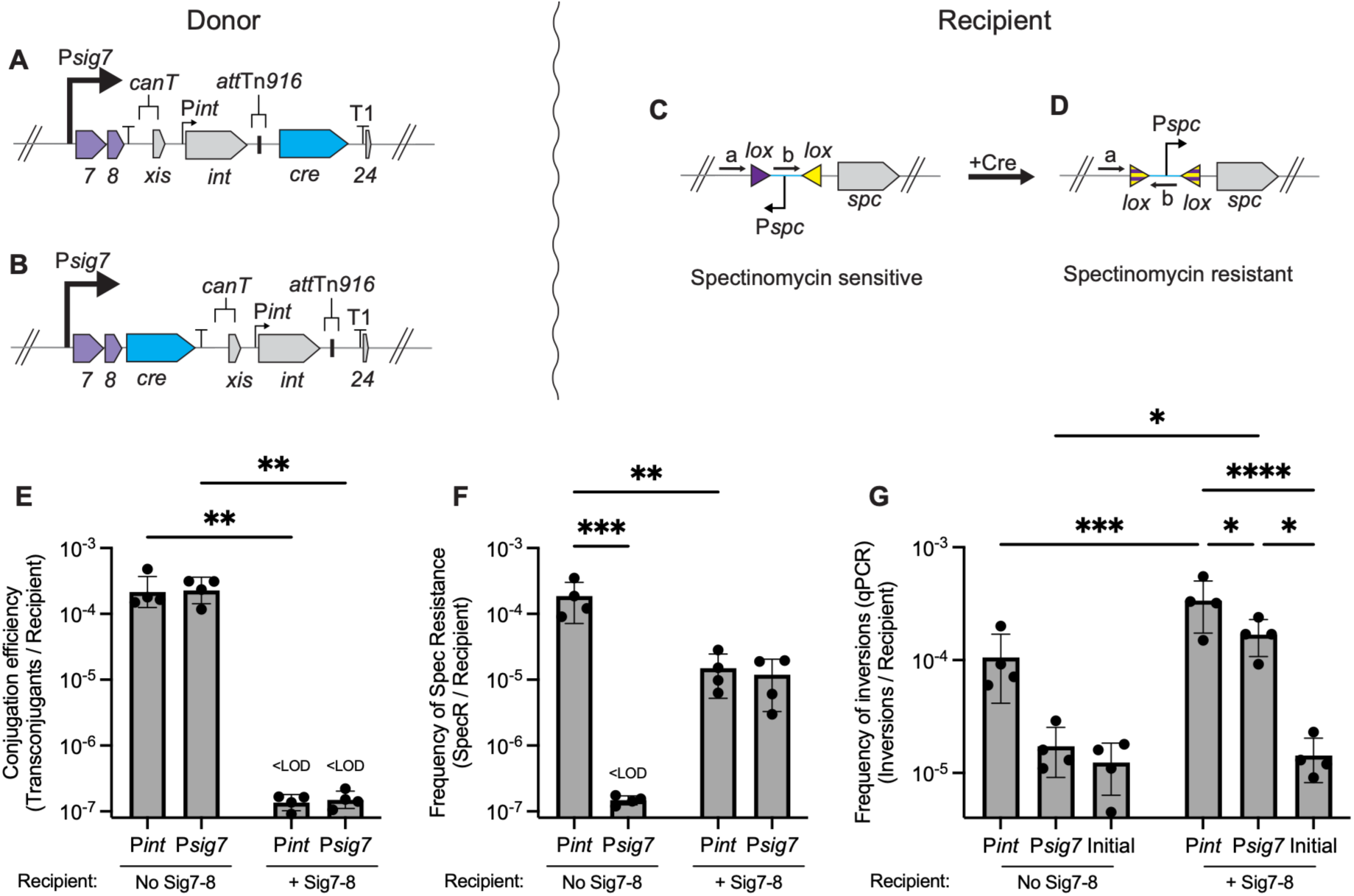
Measuring expression from P*int* and P*sig7* in transconjugants. **A-D.** Schematic of the assay for monitoring transcription from Tn*916* promoters P*int* and P*sig7* in transconjugants. The ability of the recombinase Cre to catalyze inversion between two *lox* sites was used as a reporter for transcription from a given promoter. *cre-ssrA*\* (only *cre* indicated, but all reporters contain *cre-ssrA\**) was inserted into Tn*916* downstream from the promoter of interest. The reporter containing *lox* sites is present in recipient cells. **A-B.** Cartoon of the left end of the donor Tn*916* with *cre* downstream from (**A**) P*int* (and any promoter upstream from P*int*). Note that *cre* is downstream from the attachment site, so that when the element integrates into a host chromosome, *cre* will be separated from all Tn*916* promoters. (**B**) *cre* is downstream from P*sig7-sig7-sig8* (and any promoters upstream from P*sig7*). **C-D.** Reporter for expression of *cre* that is present in recipient cells. The purple and yellow triangles depict the *lox* sites that flank the DNA fragment that can be inverted. Black arrows (labeled a and b) indicate primers used in the PCR assays to detect inversions. (**C)** Recipients prior to Cre-mediated inversion are spectinomycin sensitive as the promoter is oriented in the wrong direction. (**D**) Following Cre-mediated inversion, purple and yellow triangles mix, indicating irreversible recombination (inversion) between the *lox* sites. Inversion results in cells that are spectinomycin resistant as the promoter P*spc* can now drive transcription of *spc*. **E-G.** Results from mating the indicated donor with recipients containing the reporter for *cre* expression. Donors: P*int* indicates P*int*-*cre-ssrA\** [strain AFG835]; P*sig7* indicates P*sig7*-*cre-ssrA\** [strain AFG831]. Recipients: No Sig7-8 [strain AFG570]; +Sig7-8 [P*spank*-*sig7-8*, without IPTG; strain AFG637]. Data presented include: (**E)** conjugation efficiencies; (**F**) frequency of spectinomycin-resistance in recipients; and (**G**) Cre-catalyzed inversion frequency. Results are presented per recipient because of the use of the reporter in recipient cells. Data are from four independent experiments with error bars depicting the standard error of the mean (mean ± SEM). P-values were calculated by a two-way ANOVA comparison (*P < 0.05, **P < 0.01, ***P < 0.001; ****P < 0.0001) using the GraphPad Prism version 11. Non-significant values are not shown. **E.** Conjugation efficiencies were calculated as the number of transconjugants per input recipient. All four independent mating assays of both donors into the +Sig7-8 recipient resulted in conjugation efficiencies below the limit of detection. Matings were simultaneously conducted with donors that could not transfer (relaxase-; Δ*orf20*) which were also at the limit of detection (S2A Fig). **F.** The number of spectinomycin-resistant CFUs per input recipient was calculated. This represents the number of cells that expressed *cre* from the indicated promoter and at a level sufficient to catalyze inversion. All four independent mating assays of the P*sig7*-*cre-ssrA\** donor into the No Sig7-8 recipient resulted in no spectinomycin-resistant colonies above the limit of detection, a level similar to that seen with *Δorf20* donors (S2B Fig). **G.** The frequency of inversions per recipient was calculated as the number of Cre-mediated inversions detected by qPCR using primers ’a’ and ’b’ (panel D) per recipient. "Initial" indicates the background signal in qPCR with recipients alone (pre-mating). This was a true limit of the qPCR and not an actual background of inversions because none of the cells were resistant to spectinomycin. (See also S2C Fig for results with *Δorf20* donors.)

### Transcription from P*int* in transconjugants

We inserted *cre-ssrA\** upstream of *orf24* in the linear Tn*916*, the same location as *gfp* in Tn*916*-*gfp* (Bean et al., 2022). After excision of Tn*916*, *cre-ssrA\** is downstream of *int* and transcribed from P*int* (referred to as P*int*-*cre*) and any promoters upstream (Fig. 6A). P*int* is transcribed by the host RNA polymerase, likely containing the major sigma factor SigA (Marra et al., 1999).

As expected, we detected transcription from P*int* in transconjugants. Matings between donors containing Tn*916*-(P*int*-*cre-ssrA*\*) and recipients containing the reporter for Cre activity resulted in ∼2×10^-4^ transconjugants per recipient (Fig. 6E). Similarly, the frequency of spectinomycin resistant cells per recipient was ∼2×10^-4^ (Fig. 6F) and a similar frequency of inversions was detected by PCR (Fig. 6G). To be sure that we were not measuring separate subpopulations, one of which stably acquired Tn*916* (TetR) and another that became spectinomycin-resistant, we tested the spectinomycin-resistant colonies for resistance to tetracycline. Of 100 spectinomycin-resistant colonies, 92 were also tetracycline-resistant, indicating that they had stably acquired Tn*916*.

Conjugative transfer of Tn*916*-(P*int-cre-ssrA\**) was required to generate tetracycline-resistant transconjugants, spectinomycin-resistant cells, and inversions detected by qPCR. Using donor cells with a Tn*916* mutant that was incapable of transfer due to a deletion in the gene for the relaxase (Δ*orf20*), we detected no tetracycline-resistant transconjugants, no spectinomycin-resistant recipients, and no inversions above the background in the qPCR assay (S2 Fig). These results indicate that there was no transfer of Tn*916* DNA and no transfer of Cre-SsrA* protein from donors to recipients. Together, these results confirm that P*int* (or a promoter upstream) is expressed in transconjugants.

### P*sig7* is not expressed in most transconjugants

We inserted *cre-ssrA\** downstream of *sig8* to create a P*sig7*-*cre-ssrA*\* reporter. Transcription of *cre-ssrA\** would then be from P*sig7* and any upstream promoters (Fig. 6B) and would be regulated similarly to transcription of *sig7* and *sig8*.

As with Tn*916*-(P*int*-*cre-ssrA\**), there were ∼2×10^-4^ transconjugants per recipient using donor cells with Tn*916-*(P*sig7*-*cre-ssrA\**) (simply P*sig7*-*cre* below) (Fig. 6E). In marked contrast to donors with Tn*916-*(P*int*-*cre*), we did not detect spectinomycin-resistant cells (<10^-7^ spectinomycin-resistant cells/recipient) following mating with Tn*916*-(P*sig7*-*cre*) (Fig. 6F). These results indicate that P*sig7*-*cre* was not expressed in viable transconjugants, or was expressed at a level below that needed for Cre-mediated inversion of the reporter. We also measured the frequency of Cre-mediated inversions in recipients by qPCR, irrespective of cell viability. We detected little or no inversions from Tn*916*-(P*sig7*-*cre*) above the background in the qPCR assay (Fig. 6G). Together, these results indicate that there are few if any viable transconjugants that had transcription from P*sig7*.

### Expression of *sig7-8* in recipients caused expression from P*sig7* and prevented generation of stable transconjugants

As described above, we suspected that transcription from P*sig7* in transconjugants would limit the stable acquisition of Tn*916*. To test this, we expressed a low level of *sig7* and *sig8* in recipients. We fused *sig7-8* to P*spank* (P*spank*-*sig7-8*) and integrated this in single copy at an ectopic site (*lacA*) in the *B. subtilis* chromosome. P*spank* is leaky and there is a low level of expression, even in the absence of the inducer IPTG (Griffith and Grossman, 2008). We used this strain as recipient in a cross with donors containing Tn*916-*(P*sig7*-*cre-ssrA\**). Following mating, there were no detectable transconjugants as determined by tetracycline-resistant recipients (<10^-7^ transconjugants/recipient) (Fig. 6E). This is in marked contrast to mating into recipients that did not contain P*spank*-*sig7-8*. Tn*916*-(P*sig7*-*cre-ssrA\**) transferred to recipients, as evidenced by Cre-mediated inversions. This indicates that P*sig7* was activated in recipients by the low levels of *sig7-8* expression (Fig. 6F). The frequency of spectinomycin-resistant recipients was 1×10^-5^ (± 9×10^-6^) per recipient, but these were sensitive to tetracycline (0/82 colonies were resistant to tetracycline) indicating that they had not stably acquired Tn*916*. The frequency of Cre-mediated inversions was ∼2×10^−4^ (± 6×10^−5^) per recipient, approximately 10-fold greater than inversions into recipients without *sig7-8* (Fig. 6G). Notably, the inversion frequency was also ∼10-fold greater than the frequency of spectinomycin-resistant recipients (Fig. 6F), indicating that ∼90% of the cells that expressed *cre* from P*sig7* (detected by PCR) did not survive (survival measured by spectinomycin-resistant colonies) in a recipient expressing *sig7-8*.

Additionally, mating of Tn*916*-(P*int*-*cre*) donors into a recipient with P*spank*-*sig7-8* similarly yielded spectinomycin-resistant colonies and no tetracycline-resistant transconjugants, indicating that this donor also enters recipients with *sig7-8* but is not stably acquired (Fig. 6E, F). The frequency of spectinomycin-resistant recipients was 2×10^-5^ (± 1×10^-5^) per recipient, 10-fold less than into recipients without *sig7-8*. These were sensitive to tetracycline (0/82 colonies were resistant to tetracycline) indicating that they had also not stably acquired Tn*916* due to the presence of *sig7-8*. The frequency of Cre-mediated inversions as measured by qPCR was ∼4×10^-4^ (± 1×10^-4^) per recipient and was significantly higher than inversions into a recipient without *sig7-8* and higher than the background signal from qPCR (Fig. 6G).

Based on these results, we conclude that there are two distinct regulatory outcomes in transconjugants: 1. there is active transcription from P*int* driving expression of *int* (the recombinase gene), and 2. there is little or no active transcription from P*sig7*, at least in the majority of transconjugants. When P*sig7* is active in nascent transconjugants, there are no detectable stable transconjugants, most likely due to: 1) expression of *sig7-8* causing transcription from P*sig7* and high levels of Sig7-8 and expression of conjugation genes that cause cell death, and 2) expression of *xis* from P*sig7* preventing stable integration of the element. Thus, the inactivity of P*sig7* in nascent transconjugants enables both cell survival and stable integration of the element.

## Discussion

Previous work established that an activated Tn*916* kills its host donor cells (Bean et al., 2022), yet transconjugants survive. Our findings show that killing is due in part to high levels of Sig7-8, which can cause cell death in the absence of any other Tn*916* genes. Further, Sig7-8 are required for transcription of other Tn*916* genes that are also involved in host killing. Transconjugants survive because there is little or no expression of *sig7-8*. *sig7* and *sig8* encode a two-subunit (split) sigma factor that drives transcription from P*sig7*, the major Tn*916* promoter. In donors, this leads to high-level expression of *sig7-8* and the conjugation genes. In transconjugants, P*sig7* remains inactive due to little or no production of Sig7-8. We suspect that a relatively small fraction (∼1-3%) of nascent transconjugants express *sig7-8*, and these cells likely do not survive. If they do survive, they likely lose the element due to expression of *xis* from P*sig7*.

### Two component, split sigma factors

There are several examples of two-subunit split sigma factors, with early examples coming from bacteriophages. SP01 encodes Gp33 and Gp34, which together direct transcription of late phage genes (Losick and Pero, 2018; Tjian and Pero, 1976). Similarly, coliphage T4 encodes Gp55 and Gp33, which together comprise a two-subunit split sigma factor that activate late promoters via interaction with the sliding clamp of the phage DNA polymerase (Nechaev et al., 2004). In these bacteriophage examples, both subunits of the split sigma factors are required for transcription. Canonical sigma-70-like factors have four domains (regions 1-4). In contrast, some alternative sigma factors (group 4) contain only regions 2 and 4 (Helmann, 2002; Lonetto et al., 1992). The *B. subtilis* proteins RsoA (YvrHa) and SigO (YvrI) comprise a two-subunit split sigma factor that together have functional regions 2 and 4 (MacLellan et al., 2008). Based on results presented here, combined with previous sequence analyses, we conclude that Sig7-8 from Tn*916* functions as a two-subunit (split) sigma factor that binds to and initiates transcription from the major Tn*916* promoter P*sig7*.

### Cell death due to overproduction of alternate sigma factors

The production and/or activity of many alternate sigma factors is highly regulated (Helmann, 2010; Losick and Pero, 1981; Österberg et al., 2011; Paget, 2015; Pinto et al., 2019; Schumann, 2016). Overproduction or overactivation, whether through ectopic expression, loss of a natural repressor, or inappropriate induction, can severely alter host transcription and phenotypes. For example, sporulation sigma factors in *B. subtilis* are highly regulated and their untimely expression can disrupt vegetative growth and prevent proper sporulation (Fimlaid and Shen, 2015; Hicks and Grossman, 1996; Hilbert and Piggot, 2004; Stragier and Losick, 1996). Deleterious effects result primarily from early activation of genes needed for sporulation and perhaps partly from competition with vegetative sigma factors for limiting core RNA polymerase (Hicks and Grossman, 1996; Lord et al., 1999; Serrano et al., 2011; Sun et al., 1989).

In contrast, some alternate sigma factors appear to be toxic primarily through competition with vegetative sigma factors. There is a limiting pool of core RNA polymerase (Grigorova et al., 2006; Hicks and Grossman, 1996) and changes to levels of the major sigma or alternate sigmas can alter the proportions of different forms of RNA polymerase holoenzyme (Chelm et al., 1982). An extreme case is SigN, a sigma factor encoded on plasmid pBS32 in *B. subtilis*. SigN is normally made in response to DNA damage and accumulates through a positive feedback loop (Burton et al., 2023, 2019). When SigN is overexpressed in the absence of its regulators (encoded by some of the genes it activates), it causes rapid cell death, likely through competition with SigA (Burton et al., 2023).

Sig7-8, like SigN, is autoactivated: once sufficient Sig7-8 accumulates to activate transcription from P*sig7*, a positive feedback loop is created, driving further Sig7-8 production. This autoactivation and competition for core RNAP likely causes toxicity and explains the selective pressures to limit transcription from P*sig7* in nascent transconjugants and thereby enabling cell survival and propagation of Tn*916*.

### Autoactivating switch for Tn*916* gene expression

Sig7 and Sig8 are both required to activate transcription from P*sig7* (Celli and Trieu-Cuot, 1998). This creates a positive autoregulatory loop. Once sufficient Sig7-8 accumulates to activate P*sig7*, more Sig7-8 is produced, locking cells into a committed "on" state. This type of cell-fate commitment, often referred to as a “bistable switch” (Gardner et al., 2000; Ge et al., 2018; Süel et al., 2007), is analogous to terminal differentiation of cells with an activated ICE*clc* (Carraro et al., 2020).

Initial transcription of *sig7-8* (independent of P*sig7* and Sig7-8) can come from readthrough transcription from the upstream promoter P*tet* (Su et al., 1992). These transcripts are regulated by a transcription attenuation mechanism in which readthrough transcription increases when ribosomes stall on a small open reading frame (Su et al., 1992; Turnbough, 2019; Yanofsky, 1988). Normally, in the absence of tetracycline, there is a subpopulation of cells with enough Sig7-8 to enable transcription from P*sig7* and activate Tn*916*.

Tetracycline directly links antibiotic stress to Tn*916* activation. Tetracycline inhibits translation, causing ribosome stalling and increasing readthrough transcription into *tet*(*M*) and *sig7-8*. This increase in *sig7-8* expression results in an increase in the fraction of cells with a threshold amount of Sig7-8 and subsequent increase in activation of Tn*916*. This stochastic, infrequent activation represents a bet-hedging strategy that enables a subpopulation of cells to become Tn*916* donors and die. The proportion of bet-hedging cells increases in the presence of tetracycline.

### Interactions between Tn*916* and host cells

Tn*916* confers tetracycline resistance to its host, a clear benefit in the presence of the antibiotic. However, activation of Tn*916* causes host cell death through at least three distinct mechanisms that could be detrimental or beneficial for spread of the element, as discussed previously (Gomberg and Grossman, 2024).

At least one killing mechanism limits the spread of Tn*916*. *yqaR*, found in the defective phage *skin*, in combination with the Tn*916* genes *orf17* and *orf16*, causes killing, and deletion of *yqaR* results in a 10-fold increase in conjugation frequency (Bean et al., 2022). This indicates that host killing constrains element spread, potentially limiting the competitive advantage tetracycline resistance would provide to neighboring tetracycline-sensitive cells that lack the element (Bean et al., 2022).

This contrasts with ICE*clc* from *Pseudomonas*, where killing genes promote element transfer. That is, removal of killing genes from ICE*clc* decreases conjugation efficiency (Reinhard et al., 2013; Takano et al., 2019). ICE*clc* induces a specialized transfer-competent state that supports high levels of conjugation, thereby benefiting the element.

The role of Sig7-8-mediated killing in Tn*916* is not known. We suspect that Sig7-8 toxicity benefits the element by locking donor cells into a committed, transfer-competent state, analogous to the terminal differentiation induced by ICE*clc* (Reinhard et al., 2013; Takano et al., 2019). We also speculate that high levels of Sig7-8 outcompete other sigma factors for binding to core RNA polymerase and inhibit transcription of other genetic elements that depend on host sigma factors for gene expression. In this way, Sig7-8 might protect Tn*916* from competition with other mobile elements.

### Tn*916*-family of ICEs and positive regulation of ICE gene expression

Tn*916* is the archetype of a large, prevalent family of ICEs that all seem to have similar regulatory elements (Antezana et al., 2023; Roberts and Mullany, 2009). The key regulatory features of these elements include a major promoter that is physically separated from the conjugation genes when the element is integrated, and what we now recognize as genes for a two-subunit sigma factor that activate transcription from the main element promoter. The activator, Sig7-8, is not produced at a level sufficient for transcription from P*sig7* when the element is integrated in the chromosome. Apparently, by a stochastic mechanism, a subpopulation of Tn*916*-containing cells produce enough Sig7-8 to activate transcription from P*sig7*, creating a positive feedback loop to transcribe more *sig7-8* and the downstream *xis* and *int*, and causing Tn*916* to excise and circularize and transcribe element genes needed for conjugation. However, when Tn*916* enters a new host, the activators are absent and there is little or no transcription of element genes from the main promoter P*sig7*.

In a nascent transconjugant, activator-based regulation leads to fundamentally different outcomes than repressor-based regulation. In transconjugants, activator-based systems are de-facto inactive upon entry to a new host. In contrast, repressor-based systems are, de-facto, active upon entry as the repressor is not present. This is true of repressor-based regulation of ICEs (Auchtung et al., 2016; Bose et al., 2008; King et al., 2022; McLellan et al., 2022; Pradervand et al., 2014) and some of the best studied temperate phages (Bertani, 1953; Jacob and Wollman, 1953; Wollman et al., 1956). Repressor-based regulation is the basis for zygotic induction (Wollman et al., 1956) and immunity (repressor-mediated inhibition of acquisition of a second copy of an element) (Bertani, 1953). Entry of a repressor-regulated ICE into a new host can result in expression of conjugation genes and nascent transconjugants becoming donors prior to element repression and integration (so-called secondary transfer) (Babic et al., 2011; Bellanger et al., 2009; Johnson and Grossman, 2015).

We predict that activator-based elements, including Tn*916* and its family members, integrate into the host chromosome before the activator is made, possibly allowing them to integrate very soon after entry to a new host. Since it lacks a repressor, the presence of Tn*916* in a cell does not confer immunity against conjugative acquisition of another copy of the element (Flannagan and Clewelll, 1991; Norgren and Scott, 1991; Rice et al., 2010). We speculate that Tn*916* does not undergo secondary transfer from a nascent transconjugant. Despite clear differences in lifestyle, both activator-based and repressor-based regulatory strategies are successful given the prevalence of the Tn*916*-like family and repressor-based ICEs.

## Materials and Methods

### Media and growth conditions

*Escherichia coli* cells were grown shaking at 37°C in LB medium for routine strain constructions. *B. subtilis* cells were grown shaking at 37°C in either LB medium or MOPS (morpholinepropanesulfonic acid)-buffered S750 defined minimal medium (Jaacks et al., 1989) containing 0.1% glutamate, required amino acids (40 μg/ml phenylalanine and 40 μg/ml tryptophan), and glucose (1% w/v) as a carbon source, or on LB plates containing 1.5% agar. For most experiments, *B. subtilis* strains were grown in defined minimal medium with 1% glucose at 37°C with shaking to the indicated cell density. Where indicated, tetracycline (2.5 µg/ml) was added to Tn*916*-containing cells to increase gene expression, excision, and transfer (Celli and Trieu-Cuot, 1998; Scornec et al., 2017; Showsh and Andrews, 1992). Antibiotics were otherwise used at the following concentrations: 5 μg/ml kanamycin, 10 μg/ml tetracycline (to select for tetracycline-resistance), 100 μg/ml spectinomycin, 5 μg/ml chloramphenicol, and a combination of erythromycin at 0.5 µg/ml and lincomycin at 12.5 µg/ml to select for macrolide-lincosamide-streptogramin (MLS) resistance.

For all assays except matings, cells were first grown as light lawns on 1.5% agar plates containing 1% w/v glucose, 0.1% w/v monopotassium glutamate, and 1X Spizizen’s salts (2 g/l (NH₄)₂SO₄, 14 g/l K₂HPO₄, 6 g/l KH₂PO₄, 1 g/l Na₃-citrate·2H₂O, and 0.2 g/l MgSO₄·7H₂O) (Harwood and Cutting, 1990). Cells were resuspended from light lawns and grown at 37°C with shaking in 1X S750 defined minimal glucose medium. For mating experiments, a fresh single colony of cells was picked from an LB plate, resuspended in LB medium, and grown to exponential phase.

### Strains, alleles, and plasmids

*E. coli* strains AG1111 (MC1061 F’ *lacIq lacZM15* Tn10) and DH5α were used for plasmid construction. *Bacillus subtilis* strains were derived from JMA222, a derivative of JH642 (*trpC2 pheA1*) (Smith et al., 2014) that was cured of ICE*Bs1* (Auchtung et al., 2005) and are listed in Table 1. Strains were constructed by natural transformation (Harwood and Cutting, 1990). Unless otherwise noted, Tn*916* and derivatives are integrated between *yufK* and *yufL* (Bean et al., 2022; Wirachman and Grossman, 2024).

The following alleles were reported previously:

- Δ*comK*::*kan* (to prevent transformation into recipients during matings) (Wirachman and Grossman, 2024)
- *cgeD*::{[P*immR-immR-immA*] *kan*} expresses *immR* and *immA* and was used to repress P*xis* and allow RapI-mediated derepression (Bean et al., 2022)
- Δ*orf21* in Tn*916* (Bean et al., 2022). *orf21* encodes the coupling protein that is essential for conjugation.
- *canT*(*Δ162*) removes 162 bp between *sig8* and *xis*, leaving both genes intact and eliminating function of the transcription antiterminator (Wirachman and Grossman, 2024).
- *rpoC-his10* encodes the ß’ subunit of RNA polymerase with a fusion of 10 histidines at the C-terminus (Qi and Hulett, 1998). It was used for affinity purification of RNA polymerase and associated proteins.

We cloned P*sig7* into an ectopic site in the *B. subtilis* chromosome and used this in crosslinking and immunoprecipitation assays to determine if RNA polymerase with Sig7-8 was associated with P*sig7*. The fragment containing P*sig7* was essentially as described (Celli and Trieu-Cuot, 1998): the 5’ end is 216 bp upstream from the newly annotated *sig7* start codon (GTG) and the 3’ end is after 11 codons of *sig7*. *ycgO*::{[P*sig7*-*mapple2*] *cat*} was generated by isothermal assembly of DNA fragments containing P*sig7*, *mapple2* (McKeithen-Mead and Grossman, 2023), *cat*, and homology arms to *ycgO* and then transformation into *ycgO* in the *B. subtilis* chromosome selecting for chloramphenicol resistance.

#### Ectopic expression of *sig7* and *sig8* from ICE*Bs1* P*xis*

Constructs for ectopic expression of Tn*916* genes from the major ICE*Bs1* promoter P*xis* were made using pAFG227, a vector for integrating DNA at *amyE*. This plasmid contains a fragment of ICE*Bs1* DNA extending from *immA* to just before *helP*, including *immR* and P*xis*, cloned between the BamHI-EcoRV sites of the pDR110-derived plasmid pAJW310 (deleted for P*spank* and *lacI*). Inserts of *sig7-8* extended from the annotated transcription start site upstream of *sig7* to 59 bp downstream of the *sig8* stop codon. Specific constructs include:

- *amyE*::{[P*xis*-*sig7-sig8*] *spc*} (AFG288): wild-type *sig7-sig8*
- *amyE*::{[P*xis* empty] *spc*} (AFG289): vector control
- *amyE*::{[P*xis*-*sig7-sig8M7ns*] *spc*} (AFG308): *sig8* with a nonsense mutation at codon 7 (ATG to TAG)
- *amyE*::{[P*xis*-*sig8*] *spc*} (AFG310): *sig8* only, including the final five codons of *sig7* and the putative *sig8* ribosome binding site
- *amyE*::{[P*xis*-*sig7-flag-sig8-myc*] *spc*}: contains the same boundaries as P*xis sig7-sig8* with epitope tags at the 3’ ends—for Sig7, a GGGGS linker and FLAG epitope (*sig7-flag*); for Sig8, a GSA linker and 3x-Myc tag (*sig8-myc*), with the putative *sig8* ribosome binding site preserved. This construct was used for co-immunoprecipitation and chromatin immunoprecipitation assays.

#### P*spank*-inducible clones

The LacI-repressible-IPTG-inducible promoter P*spank* was used to control expression of some genes. Constructs included:

- *lacA*::{[P*spank*-*rapI*] *tet*} (strain SAM497) contains *rapI* from ICE*Bs1* cloned downstream of the LacI-repressible-IPTG-inducible promoter P*spank* and integrated into the nonessential gene *lacA*. Addition of 1 mM IPTG induces *rapI* expression, leading to RapI-mediated inactivation of the repressor ImmR, which allows derepression of P*xis*. A DNA fragment extending from 29 bp upstream of the *rapI* start codon to 15 bp downstream of the stop codon was PCR-amplified and cloned downstream of P*spank* (in plasmid pSAM487), then integrated into *lacA* in JMA222 by transformation, selecting for tetracycline resistance.
- *•lacA*::{[P*spank*-*sig7-sig8*] *mls*}: *sig7* and *sig8* from Tn*916*, using the same boundaries as described for P*xis*-*sig7-sig8*, were placed under control of P*spank* at *lacA*. P*spank* and *lacI* were amplified from pDR110. *mls* was cloned from ESW271 (Wirachman and Grossman, 2024). DNA was assembled by isothermal assembly (Gibson et al., 2009) and transformed into *B. subtilis* selecting for MLS resistance.
- *•amyE*::{[P*spank*-(*Ec*)*sspB*] *mls*}: The *E. coli*-derived *sspB/ssrA*\* system that was adapted for targeted protein degradation in *B. subtilis* was used essentially as described (Griffith and Grossman, 2008). Production of the adaptor protein SspB (from P*spank*-*sspB*) enables Clp-mediated degradation of Cre-SsrA*. A derivative of the previously described *amyE*::{[P*spank*-*sspB*] *spc*} was generated with *mls* replacing *spc* by PCR assembly and isothermal assembly. DNA was transformed into *B. subtilis* selecting for MLS resistance; transformants were MLS-resistant and spectinomycin-sensitive.

#### Unmarked deletions and mutations in Tn*916*

Unmarked deletions, insertions, and mutations in Tn*916* were generated by allelic replacement using the following approach: flanking homology regions were amplified for each deletion and mutation and inserted by isothermal assembly (Gibson et al., 2009) into the EcoRI and BamHI sites of pCAL1422 (a plasmid containing *E. coli lacZ* and *B. subtilis cat*), as previously described (Thomas et al., 2013). Resulting plasmids were used to transform appropriate *B. subtilis* strains, selecting for integration by single crossover recombination. Transformants were passaged and screened for loss of *lacZ* and chloramphenicol sensitivity, and checked by PCR and sequencing. Specific mutations include:

- Δ*orf20*: deletes 306 codons of *orf20*, from codon 90 to the stop codon, preserving the origin of transfer *oriT*
- Δ*xis*: deletes 40 codons of *xis*, from codon 26 to the final two codons
- Δ*orf*(*19-13*): retains 37 bp following the *orf20* stop codon and the final 27 bp of *orf13*, deletes the intervening six open reading frames
- *•sig7M4I* changes the ATG (methionine) that we believe is codon 4 to ATA (isoleucine) (S1A Fig). Tn*916* with this mutation has a wild type phenotype (S1 Fig), indicating that either the ATA functions normally as a start codon (unlikely) or that the actual start codon is upstream, most likely the GTG three codons upstream. This ATG was previously predicted to be the start codon of *orf7.2* (Su and Clewell, 1993)
- *sig7M4ns* changes the ATG (methionine) that we believe is codon 4 to TGA (nonsense) (S1A Fig)
- *•sig8M1I*: changes the predicted start codon from ATG to ATA (methionine to isoleucine) while preserving the stop codon for *sig7* (S1B, C Fig)
- *•sig8M7ns*: changes the seventh codon from ATG (methionine) to TAG (nonsense)

#### *cre* reporters for monitoring gene expression in transconjugants

Tn*916*-(P*sig7*-*cre-ssrA*\*) contains *cre-ssrA*\* downstream from *sig8*. Sequences with a ribosome binding site (5’ AAGGAGAACGCAT 3’), *cre* from pDR244 (Koo et al., 2017), and the *ssrA*\* tag (Griffith and Grossman, 2008) were assembled and inserted directly downstream of the *sig8* stop codon.

Tn*916*-(P*int*-*cre-ssrA*\*) contains the same *cre-ssrA*\* construct inserted into Tn*916* 29 bp upstream of the *orf24* start codon, in the same location as *gfp* in Tn*916-gfp* previously described (Bean et al., 2022). In this location, *cre-ssrA*\* is not expressed when Tn*916* is integrated in the chromosome (as long as there is no host promoter upstream); upon excision, *cre-ssrA*\* is expressed from P*int* and upstream Tn*916* promoters.

### RpoC-His10 affinity purification

Indicated strains containing *rpoC-his10* and *amyE*::{[P*xis*-*sig7-flag-sig8-myc*] *spc*} were grown as described above (Media and growth conditions). Expression was induced by addition of 1 mM IPTG (inducing P*spank*-*rapI*) for one hour, and cells were harvested in mid-exponential phase (OD₆₀₀ ∼0.3–0.8). Cells were pelleted, resuspended in 10 ml of phosphate-buffered saline (PBS), pelleted again, and frozen at −80°C. Pellets were resuspended in 450 µl of resuspension buffer (20% sucrose, 50 mM NaCl, 10 mM Tris-HCl pH 8.0) supplemented with 10 mg/ml lysozyme, 10 µl of EDTA-free protease inhibitor cocktail (P8894; Sigma-Aldrich), and 250 units of Benzonase nuclease, and incubated at 37°C for 30 minutes. Lysis was completed by addition of 450 µl of lysis buffer (100 mM Tris-HCl pH 7.0, 300 mM NaCl, 1% Triton X-100). Lysates were spun at 4°C for five minutes, and an aliquot was saved as an untreated input sample. The remaining clarified lysate was incubated with TALON Metal Affinity Resin (Takara Bio) according to manufacturer protocols. Resin was washed with a 1:1 mixture of resuspension buffer and lysis buffer, centrifuged, and supernatant was gently removed using bent-needle aspiration. Bound protein was eluted with 300 mM imidazole in a 1:1 mixture of resuspension buffer and lysis buffer.

Fractions (lysate, unbound, wash, eluted) were separated on 15% SDS-PAGE and transferred to 0.2 µm nitrocellulose membrane using the Trans-Blot SD Semi-Dry transfer cell (Bio-Rad). Blots were blocked for one hour with Odyssey Blocking Buffer. Mouse M2 anti-FLAG (1:10,000; Millipore Sigma) or mouse anti-c-Myc (1:2,000; Thermo Fisher) primary antibodies were used, followed by five 5-minute washes with PBST (PBS + 0.2% Tween). Goat anti-mouse antibodies with conjugated fluorophore (1:10,000; Avantor/VWR) were used as secondary antibody. Blots were imaged on a LI-COR scanner.

### Crosslinking and immunoprecipitation

Cells containing *ycgO*::{[P*sig7*-*mapple2*] *cat*} and *amyE*::{[P*xis*-*sig7-flag-sig8-myc*] *spc*} were grown and expression from P*xis* was induced as described for RpoC-His affinity purification above. Cells were crosslinked with 1% formaldehyde for 5 minutes at room temperature and quenched with 0.25 M glycine for 5 minutes while gently rocking. Cells were pelleted, washed, and resuspended in 1.55 ml of 2x resuspension buffer (20% sucrose, 50 mM NaCl, 10 mM Tris-HCl pH 8.0) supplemented with 1 mg/ml lysozyme and 16 µl of protease inhibitor cocktail (P8894; as above), incubated at 37°C for 30 minutes, and lysed with 1.55 ml of lysis buffer as described above. DNA was sheared by sonication (one minute at 15% amplitude; 0.3" on, 0.3" off). Insoluble debris was removed by centrifugation and the supernatant was transferred to a fresh tube. An aliquot of 40 μl ("total" DNA control) was removed and saved. Protein-DNA complexes were immunoprecipitated for two hours at room temperature using end-over-end rotation with mouse M2 anti-FLAG (1:1,000; Millipore Sigma) or mouse anti-c-Myc (1:200; Thermo Fischer), followed by incubation with 30 µl of 50% protein A-Sepharose slurry for one hour at 4°C. Complexes were collected, washed six times with 1x IP buffer (resuspension buffer) and once with TE (10 mM Tris [pH 8], 0.1 mM EDTA). The slurry was resuspended in 100 µl of elution buffer (50 mM Tris [pH 8], 10 mM EDTA, 1% SDS), incubated at 65°C for 10 minutes (first elution), and a second elution was conducted using TE/SDS buffer (10 mM Tris [pH 8], 1 mM EDTA, 0.67% SDS). Samples and "total" DNA control were de-crosslinked overnight at 65°C, treated with 250 µl of TE and proteinase K (20 µl/ml) for 2 hours at 37°C, and DNA was purified using QiaQuick columns.

### qPCR for chromatin immunoprecipitation

qPCR was performed using SsoAdvanced SYBR master mix and CFX96 Touch Real-Time PCR system (Bio-Rad), quantified using the standard curve method. Standard curves were generated using *B. subtilis* genomic DNA containing wild-type Tn*916*. Primers to quantify P*sig7*: oAFG264 (5’-GTACGTCCACCAATGTGGTTT-3’) and oAFG255 (5’-TCCCAATGTGAAAGGGGGAT-3’). Primers to quantify the negative control locus *cah*: oAFG267 (5’-CAACTTCGAACGGGCCATTG-3’) and oAFG268 (5’-TGTCTTCATCGCCTGCACTT-3’). Relative P*sig7* abundance (Cq values) was normalized to total genomic DNA.

### qPCR to measure Tn*916* excision

qPCR was used to monitor excision (and therefore activation), essentially as described previously (Bean et al., 2022; Wirachman and Grossman, 2024; Wright and Grossman, 2016). Briefly, genomic DNA was isolated using the Qiagen DNeasy kit with 40 mg/ml lysozyme. qPCR was performed using SsoAdvanced SYBR master mix and CFX96 Touch Real-Time PCR system (Bio-Rad), quantified using the Pfaffl method (Pfaffl, 2001). Standard curves were generated using *B. subtilis* genomic DNA containing an empty Tn*916* chromosomal attachment site (*att1*) and the nearby locus *mrpG*.

Excision frequencies were calculated as the number of copies of *att1* divided by the number of copies of *mrpG*. We used previously described primers (LDW): oLW542 (5’-GCAATGCGAT TAATACAACG ATAC-3’) and oLW543 (5’-TCGAGCATTC CATCATACAT TC-3’) to amplify the empty chromosomal attachment site (*att1*). oLW544 (5’-CCTGCTTGGG ATTCTCTTTA TC-3’) and oLW545 (5’-GTCATCTTGC ACACTTCTCT C-3’) amplified a region within the nearby locus, *mrpG*.

### Growth and viability assays

Strains containing P*xis*-*sig7-sig8* were grown in defined minimal medium with 1% glucose to early exponential phase (OD₆₀₀ = 0.1). 1 mM IPTG was added to induce P*spank*-*rapI*, causing derepression of P*xis*. Relative viability was calculated as CFUs/ml following induction (expressing *sig7* and *sig8*) divided by the CFUs/ml prior to induction with IPTG.

### Mating and conjugation assays

#### Standard Tn*916* mating assay

Mating assays were performed essentially as described previously (Bean et al., 2022). Briefly, donor strains containing Tn*916* (tetracycline-resistant) or derivatives were grown in LB medium to early exponential phase (OD₆₀₀ ∼0.2). Tn*916* activation was stimulated with 2.5 μg/ml tetracycline for one hour. Donor and recipient cells (5 total ODs) were mixed in a 1:1 ratio, filtered, and mating filters were washed with LB and placed on 1X Spizizen’s salts 1.5% agar plates at 37°C for 2.5 hours. Cells were harvested and plated on selective media. CFUs of donors (tetracycline-resistant), recipients (chloramphenicol- or kanamycin-resistant), and transconjugants (tetracycline/chloramphenicol- or tetracycline/kanamycin-resistant) were determined pre- and post-mating. Conjugation efficiency was calculated as the CFUs of transconjugants per donor at the end of mating.

#### Tn*916-cre*-reporter mating assay

Mating assays were performed essentially as described above. Strains with Tn*916-*P*sig7-cre-ssrA\** (AFG831) and Tn*916*-P*int-cre-ssrA\** (AFG835) were used as donors. P*spank*-*sspB* (in the donors) was induced with 1 mM IPTG two hours before mating. Conjugation efficiency was calculated as the frequency of CFUs of transconjugants per recipient at the end of mating. Following mating and before plating for CFUs, a sample of cells was added to ice-cold methanol, pelleted, and stored for DNA extraction.

The reporter for inversion was present in recipients and included *spc* with an upstream promoter in the opposite orientation flanked by *lox* sites (ELC69). Cre-mediated inversion between the *lox* sites results in an active promoter transcribing *spc* and viable transconjugants with the inversion are resistant to spectinomycin. The inversion can also be detected by PCR using DNA from all cells, both viable and non-viable. Thus, the inversion frequency of the Cre-reporter present in recipients was determined by plating on LB with spectinomycin and by qPCR (McKeithen-Mead and Grossman, 2023) using previously described primers oSAM776 (5’-CCAGTCACGTTACGTTATTAGTTATAG-3’) and oSAM777 (5’-TACCGCACAGATGCGTAAG-3’) to quantify inversions (*spc*), and oLW104 (5’-GCGACGGAGAGTTAGGTTATTGG-3’) and oLW107 (5’-TTGAAGTCATTCTTTACAGGAGTCC-3’) to quantify the control locus (*cat*). Standard curves were generated using *B. subtilis* genomic DNA from a strain with the inverted *lox-spc-lox* cassette.

## Acknowledgments

We thank current and former lab members for sharing ideas, strains, and reagents, and Erika Wirachman and Mary Anderson for comments on the manuscript. Generative AI was used for minor editing and proof-reading.

Research reported here is based upon work supported, in part, by the National Institute of General Medical Sciences of the National Institutes of Health under award numbers R35 GM148343 to ADG. ASFG was supported, in part, by funds from the MIT Schimmel Family Program in Life Sciences and an MIT School of Science Community Service Fellowship. Any opinions, findings, and conclusions or recommendations expressed in this report are those of the authors and do not necessarily reflect the views of the National Institutes of Health, nor those of any other funder.

---

**S1 Figure.**
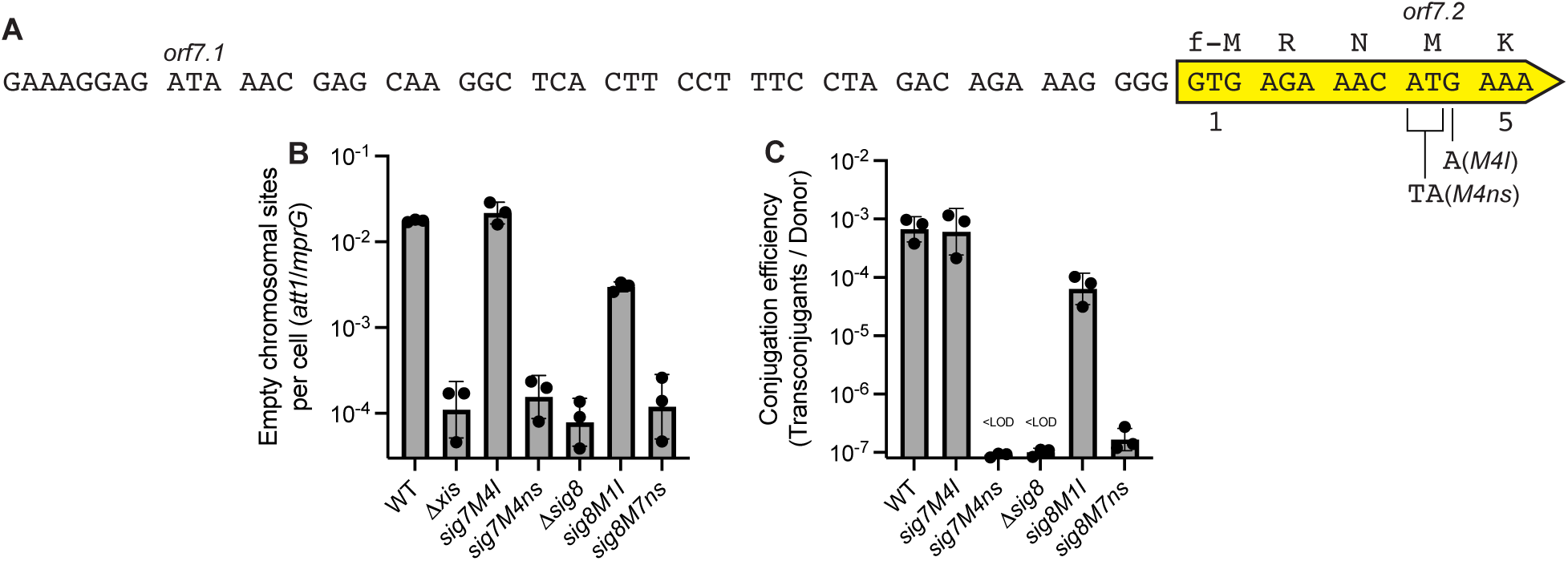
Assessing possible start codons for *sig7* and the requirement for *sig7-8* in excision and conjugation. **A.** The DNA and predicted protein sequence of the beginning of the *sig7* open reading frame. The previously proposed open reading frames *orf7.1* and *orf7.2* are indicated above the sequence and above the previously proposed start codons, ATA and ATG, respectively. Use of the ATA codon proposed as the start codon for *orf7.1* seems unlikely, although there a small number of examples of ATA start codons in bacteria have been reported (Hecht et al., 2017; Kohl et al., 2026; Peijnenburg et al., 1990; Romero and García, 1991). Our results indicate that the ATG proposed as the start codon for *orf7.2* is unlikely to function as the start codon unless ATA created by the M4I (ATG to ATA) mutation can also serve as a start codon. The new proposed GTG start codon is numbered 1 and indicated with f-M above the DNA sequence. Mutations are indicated below the sequence and include M4ns (ATG to TGA) and M4I (ATG to ATA). **B-C.** Excision (B) and conjugation (C) frequencies are presented for the indicated strains and were determined as described in Figure 3. Strains included: wild-type Tn*916* [WT, strain AFG71]; Tn*916 Δxis* [strain AFG221]; Tn*916* with a missense mutation in what we now predict to be the fourth codon of *sig7*, [*sig7M4I*; strain AFG699]; Tn*916* with a nonsense mutation in *sig7* [*sig7M4ns*; strain AFG698], Tn*916* with a deletion in *sig8* [*Δsig8*; strain AFG457]; Tn*916* with a missense mutation in the annotated start codon of *sig8* [*sig8M1I*; strain AFG454], and Tn*916* with a nonsense mutation in the seventh codon of *sig8* [*sig8M7ns*; strain AFG455]. Data presented are from independent experiments with error bars depicting the standard error of the mean (mean ± SEM). Data for WT, *Δxis*, *sig7*(*M4ns*) and *sig8*(*M7ns*) are the same as in Figure 3 and are shown here for comparison. **B.** Excision frequency was determined by measuring the amount of DNA from an ’empty’ integration site (*yufKL*), indicative of Tn*916* excision compared to the amount of DNA from a nearby site (*mrpG*). Tn*916 Δxis* [strain AFG221] is incapable of excision and was used as a control to determine the limit of detection in the qPCR assay. **C.** Conjugation efficiencies were calculated from mating the indicated donor (above and indicated below each bar) with a kanamycin-resistant recipient [strain ELC37], and measuring the number of transconjugants (resistant to tetracycline and kanamycin) per initial donor.

**S2 Figure.**
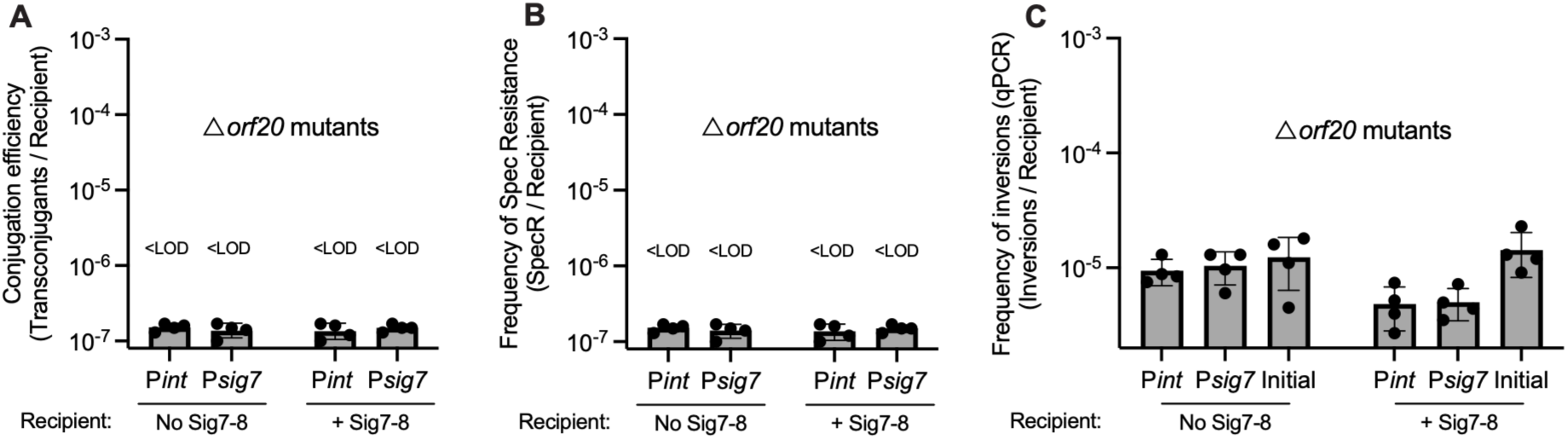
Transfer of Tn*916* is required for Cre-mediated inversions. Strains containing Tn*916 Δorf20* (relaxase mutant, incapable of conjugative transfer) and the indicated allele were crossed with the indicated recipient that contained the reporter for *cre* expression. Tn*916*-containing strains included Tn*916* P*int*-*cre-ssrA\** Δ*orf20* [P*int*, AFG833] and Tn*916* P*sig7*-*cre-ssrA\** Δ*orf20* [P*sig7*, AFG829]. Recipient strains: without Sig7-8 [No Sig7-8, AFG570] and with Sig7-8 [+Sig7-8, AFG637]. Data presented are from independent experiments with error bars depicting the standard error of the mean (mean ± SEM). Efficiencies were calculated as described in Figure 6. **A**) Conjugation efficiencies. **B**) Frequency of spectinomycin-resistance in recipients (activated promoters). **C**) Inversions per recipient.

